# Temperature and pH-dependent Potassium Currents of Muscles of the Stomatogastric Nervous System of the Crab, *Cancer borealis*

**DOI:** 10.1101/2025.10.06.680680

**Authors:** Kathleen Jacquerie, Ani Poghosyan, David J. Schulz, Eve Marder

**Affiliations:** Volen Center and Biology Department, Brandeis University, Waltham, MA 02454; Biology Department, University of Missouri, Columbia, Missouri

**Keywords:** neuromuscular junctions, 2-pore channels, crustaceans, central generators, pyloric rhythm, gastric mill rhythm, Atlantic Ocean temperature

## Abstract

Marine crustaceans, such as the crab *Cancer borealis,* experience large fluctuations in temperature and pH, yet their stomatogastric neuromuscular system must remain functional to support feeding. We examined 16 of the ∼40 pairs of stomach muscles and found that warming consistently hyperpolarized muscle fibers (∼10 mV per 10 °C) and reduced excitatory junctional potentials and currents. Voltage-clamp analysis in the gastric muscle 5b (gm5b) revealed a temperature-activated conductance with a reversal potential near the potassium reversal potential, consistent with a potassium current, and insensitive to tetraethylammonium. Quantitative RT-PCR identified expression of two putative two-pore domain potassium (K2P) channels in these muscles.

Muscle responses were also strongly influenced by extracellular pH. We observed an optimal operating window between pH 6.7–8.8; outside this range, responses diminished and abnormal activity, including spontaneous firing, appeared. Voltage-clamp recordings confirmed pH modulation of the same potassium conductance. Together, these results demonstrate that muscle excitability in *Cancer borealis* is shaped by temperature- and pH-sensitive currents, presumably carried by K2P channels.

Functionally, these channels provide a plausible mechanism for stabilizing neuromuscular output despite environmental perturbations. As temperature increases, the pyloric and gastric rhythms accelerate, increasing synaptic drive to muscles. Activation of K2P channels counterbalances this input by reducing excitability, thereby preventing over-contraction and extending its dynamic range. This work highlights a muscle-intrinsic contribution to the well-known robustness of the stomatogastric system and identifies K2P channels as key players in adapting motor performance to changing environments.

**Highlights:**

- *Cancer borealis* stomach muscles are sensitive to temperature and pH.
- Warming hyperpolarizes fibers and reduces synaptic response amplitude.
- *KCNK1* and *KCNK2*, K2P channels mediate temperature- and pH-dependent conductances.
- Muscle K2Ps lower excitability, extending the functional range at high temperatures.

**In Brief:** The crab *Cancer borealis* experiences large fluctuations in temperature and pH. We show that its stomach muscles hyperpolarize with warming and operate optimally within a narrow pH window. These effects are mediated by K2P channels (*KCNK1, KCNK2*), which reduce excitability and help maintain robust neuromuscular function under environmental change.

**Graphical abstract:** 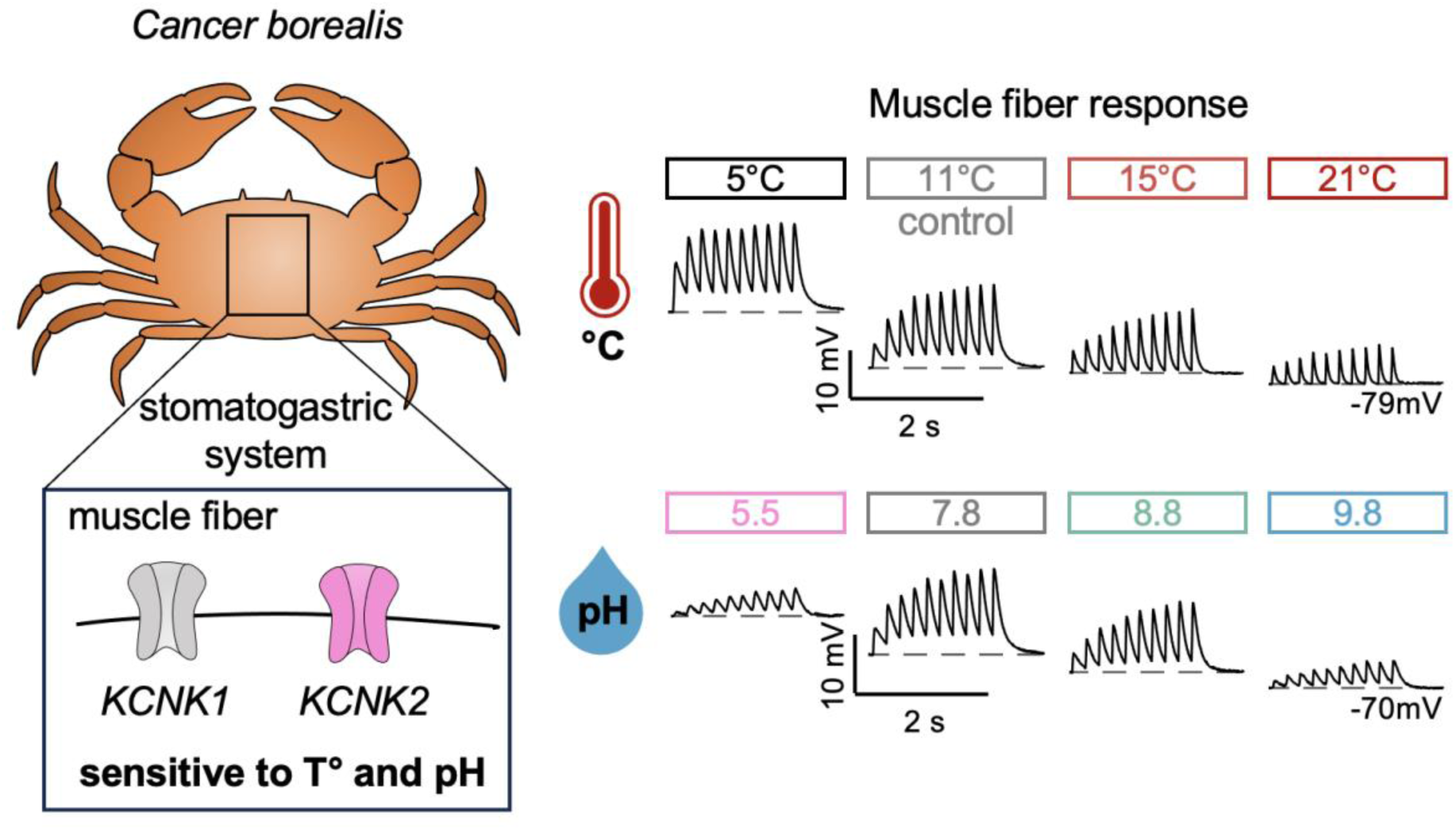

## Introduction

Many long-lived marine crustaceans, such as lobsters and crabs, face daily, monthly, and seasonal changes in water temperature and other environmental conditions, such as pH and salinity. Specifically, in the New England Atlantic Ocean, the environment in which the crab, *Cancer borealis*, is found, water temperatures vary from ∼3 °C to ∼ 25 °C [1-4]. Nonetheless, the function of these animals must be preserved, despite the strong influence that temperature has on all biological processes, including electrical excitability and synaptic transmission.

The crustacean foregut is a complex mechanical structure, involving the coordinated movements of more than 40 pairs of striated muscles, innervated by ∼20 excitatory motor neurons found in the stomatogastric ganglion (STG) [5]. Many of the STG motor neurons provide the sole innervation of one or more muscles of the stomach, using either glutamate or acetylcholine to excite the muscles [6-8]. In this study, we take advantage of the simple innervation patterns of the stomatogastric neuromuscular system to study the effects of temperature and pH on these muscles.

It has long been known that crustacean muscles hyperpolarize when they are warmed, but this was conventionally attributed to effects on the Na^+^/K^+^ pump and/or changes in leak conductance [9-12]. Understanding the biophysical mechanisms underlying this hyperpolarization and their potential functional consequences is one of the goals of this work.

Recent work has characterized a family of two pore K^+^ channels, which are either temperature and/or pH-sensitive [13-16]. In humans and mice, some of these channels are implicated in mechanosensation, temperature sensing, pain sensing, and are modulated by general aesthetics or bioactive molecules such as arachidonic acid [17-19]. In vertebrates, there are approximately 15 genes that encode this family of channels [20]. In *Drosophila,* there are approximately 10 putative homologs [21], while in *C. elegans,* about 50 have been tentatively identified [20]. Recent studies of the *Homarus americanus* [22] and *Cancer borealis* [23] genomes have identified only two candidate 2-Pore K^+^ channel candidates thus far, which we have studied here.

We now present evidence that suggests that the effects of temperature and/or pH on the resting membrane potential of crustacean muscles are mediated by activation of 2-Pore K^+^ channels. We suggest that the expression of these 2-Pore K^+^ channels might increase the dynamic range of frequencies over which the stomach can operate as a consequence of changes in environmental temperature.

## Results

Figure 1A shows a diagram of the stomach muscles of *Cancer borealis* studied here. These include the cpv1ab muscles that are innervated by the cholinergic Pyloric Dilator (PD) motor neurons, and muscles that are innervated by glutamatergic motor neurons Lateral Gastric (LG) (gm5b; gm6; gm8a,b), Dorsal Gastric (DG) (gm4b; gm4c), Inferior Cardiac (IC) (cv2; gm5a), Lateral Pyloric (LP) (p1; cpv4; cpv6), Pyloric (PY) (p2; p8), and Lateral Posterior Gastric (LPG) (p7). Figure 1B is a schematic showing the recording configuration used to stimulate the innervating motor nerves with a suction electrode while recording intracellularly from the muscle fibers.

**Fig. 1.**
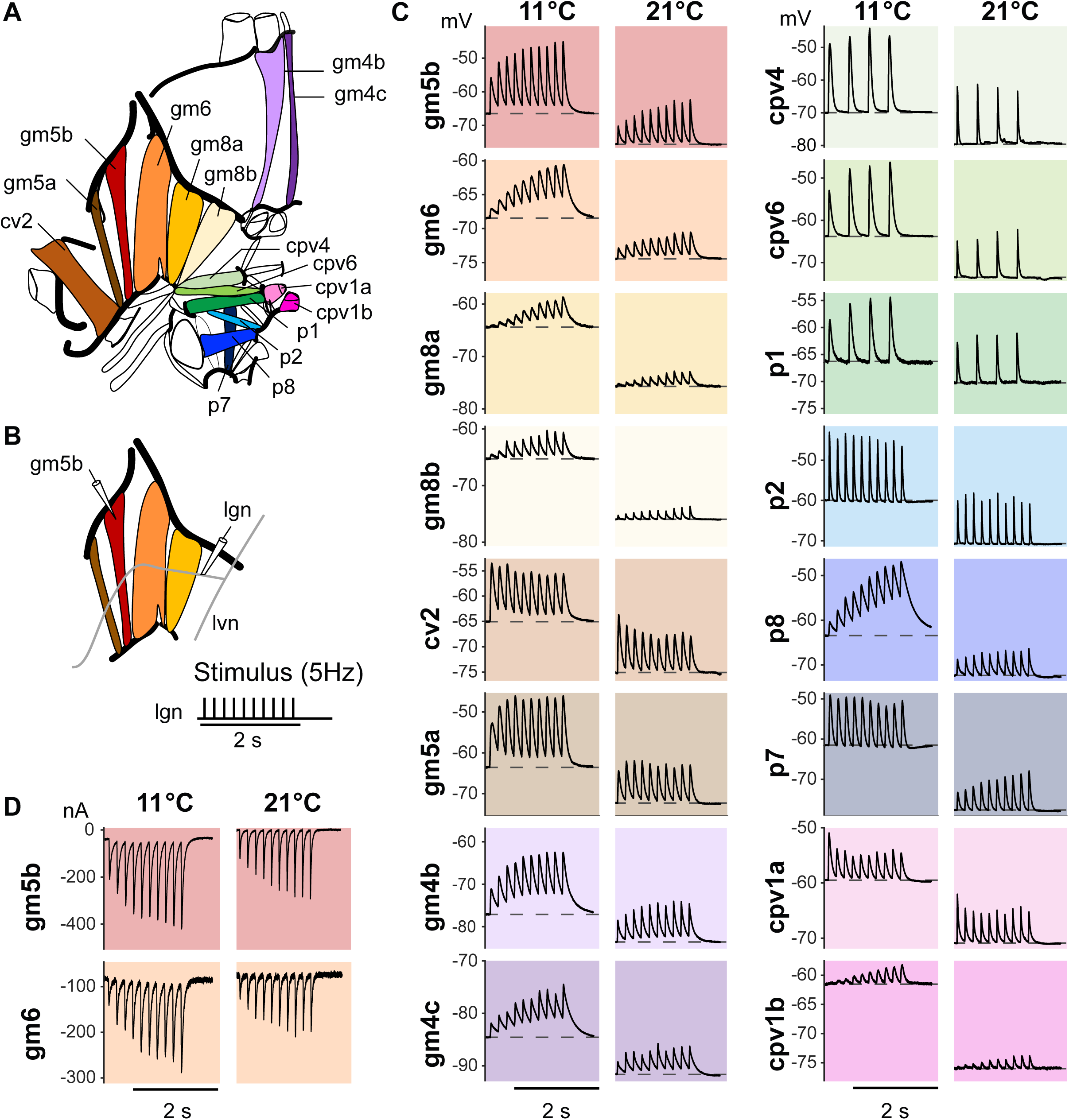
A. Schematic of the stomatogastric musculature based on the nomenclature from [5], including gastric muscles (gm), pyloric muscles (p), and cardio-pyloric (cpv) muscles—adapted from [47]. Only the left side is shown, with the anterior directed toward the bottom. B. Schematic of the experimental setup, showing in vitro dissected muscles with the innervating nerves attached. The LG neuron projects its axon to the lateral ventricular nerve (lvn) and the lateral gastric nerve (lgn). The nerve lgn is stimulated via a suction electrode with a 2-second train at 2 or 5 Hz. The EJPs are recorded via an intracellular electrode in the muscle fiber. The same setup is used to record any muscle fiber with the associated innervating nerve. C. Effect of temperature on the EJPs and resting membrane voltage (dashed gray line). Increasing temperature leads to hyperpolarization and a decrease of EJP amplitude. D. Effect of temperature on nerve-evoked EJC. A gm5b and gm6 muscle fiber are clamped in a two-electrode voltage-clamp configuration, where the membrane is clamped at -80mV. The lateral gastric nerve (lgn) is stimulated at 5 Hz for 2 seconds at 11°C and 21°C. The EJC decreases with increasing temperature.

Figure 1C illustrates the effect of increasing the saline temperature from 11^°^C to 21^°^C on the amplitude of the evoked Excitatory Junctional Potentials (EJPs) and the resting membrane potential in examples of 16 of the more than 40 pairs of stomach muscles. While the EJPS recorded vary in starting amplitude and in the extent to which they show facilitation and depression, in all recordings, as temperature was increased, EJP amplitude decreased and the membrane potential hyperpolarized. When gm5b and gm6 fibers were voltage-clamped at -80mV (Fig. 1D), the resultant Excitatory Junctional Currents (EJCs) also decreased in amplitude at 21^°^C, while retaining their characteristic facilitation.

The hyperpolarization resulting from increased temperature was seen in recordings virtually all isolated muscle fiber, and was characteristically ∼10mV with a temperature increase of 10^°^C. This occurred regardless of the innervating neuron or its neurotransmitter. The generality of this result allowed us to further quantify it and study its mechanism in a smaller subset of muscles.

Figure 2A shows data from more than 164 muscle fibers from 6 different muscles, including those innervated by the LG, PD, and LP neurons, in response to temperature increase. We voltage-clamped fibers from 3 muscles, gm5b, gm6, and cpv1a (Fig. 2B), which showed that the median input resistance of muscle fibers was modestly decreased (gm5b, -37%; gm6, -43%; cpv1a, -27%) by temperature elevation. Nonetheless, all of the changes in EJP and EJC amplitude are unlikely to be accounted solely by the changes in input resistance alone, as in some muscles, the rate of facilitation and depression was altered, thus affecting the envelope of membrane depolarization. Moreover, hyperpolarization alone would increase EJP amplitude by increasing the driving force on the transmitter-activated conductances.

**Fig. 2.**
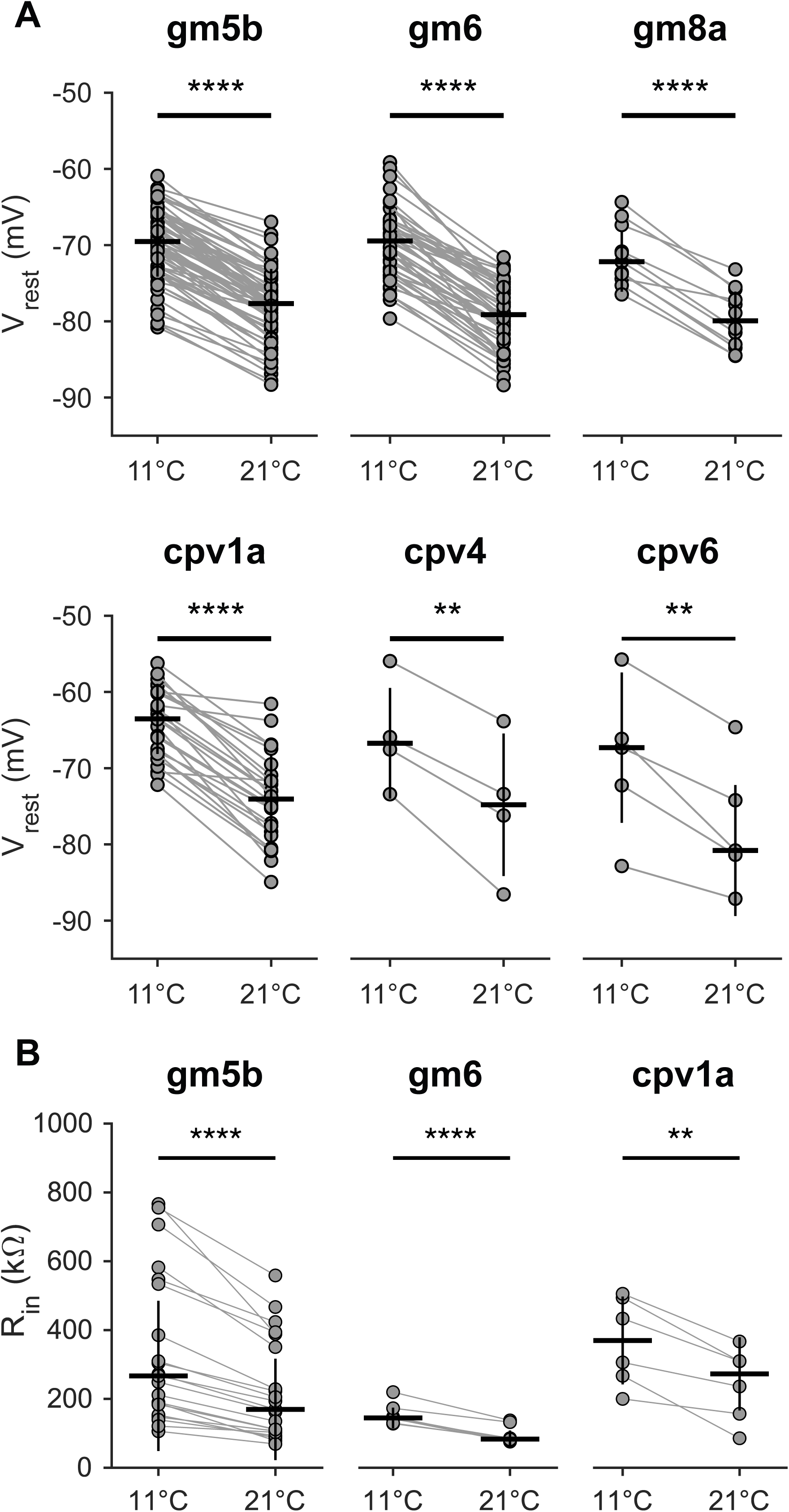
A. Quantification of the effect of temperature on the resting membrane voltage (V_rest_) for LG muscles (gm5b, gm6, gm8a), PD muscle (cpv1a), and LP muscle (cpv4, cpv6). Horizontal and vertical lines represent the median and standard deviation, respectively. Individual points correspond to single muscle fibers. Paired t-tests were performed between 11°C and 21°C (gm5b: n=68, p = 0.00000; gm6: n=50, p = 0.00000; gm8a: n=11, p = 0.00000; cpv1a: n=26, p = 0.00000; cpv4: n=4, p = 0.0057; cpv6: n=5, p = 0.0068. 0.00000 indicates p<1e^-6^). B. Quantification of the effect of temperature on the input resistance (R_in_) for LG muscles (gm5b, gm6) and PD muscle (cpv1a). R_in_ is measured in a two-electrode voltage clamp by recording the current at different voltage steps. R_in_ decreases in all three muscles. The horizontal and vertical lines represent the median and standard deviation, respectively. Each dot represents a single muscle fiber. Paired t-test is performed (gm5b: n=20, p = = 0.00000; gm6: n=8, p = 8.05e^-6^; cpv1a: n=6, p = 0.0033).

To determine the probable conductance responsible for the temperature-dependent change in the resting membrane potential (V_m_), we voltage-clamped fibers at 11^0^C and 21^0^C (Fig. 3A) and plotted them overlaid (Fig. 3B). The intersection between these curves is the reversal potential (E_rev_) of the current that is affected by temperature. For the recording shown in Figure 3B, in normal saline, E_rev_ was -86 mV, which suggests that an increase in K^+^ conductance could be responsible. To determine if this were the case, we reexamined E_rev_ in low (0.5K^+^) and high (2.K^+^) to shift the K^+^ equilibrium potential (E_K_). Figure 3B shows that, as predicted, E_rev_ of the temperature-activated current hyperpolarized in 0.5K^+^, and depolarized in 2.0K^+^. Pooled data from many fibers are shown for these manipulations in Figure 3C, and show shifts in the reversal potential of the temperature-induced current consistent with the activation of a K^+^ current.

**Fig. 3.**
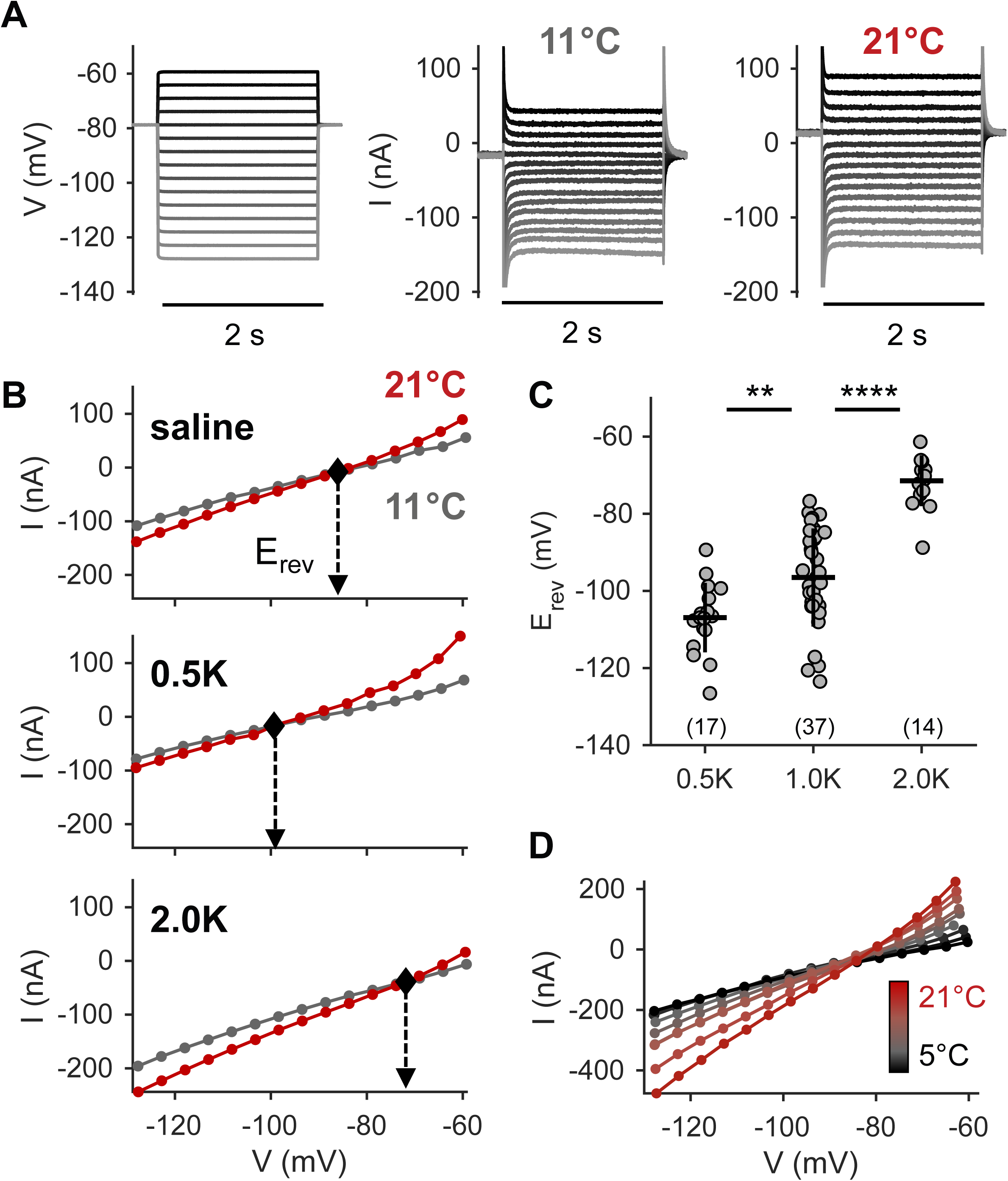
Voltage clamp assessment of the temperature sensitivity of the channel in gm5b. A. Example current traces acquired while stepping the membrane potential (from -130 mV to -60 mV over 2 seconds, left) at 11°C (center) and 21°C (right). B. IV curve of the muscle fiber at 11°C and 21°C in saline (top), marked with the reversal potential (E_rev_, diamond), which is shifted in low K^+^ (0.5xK, center) and high K+ (2xK, bottom). C. The shift in E_rev_ follows the Nernst equation, revealing a potassium channel. The horizontal and vertical lines represent the median and standard deviation, respectively. Each dot represents a single muscle fiber. (Wilcoxon rank-sum test; 0.5xK - 1.0xK: p = 0.002; 1.0xK - 2.0xK: p = 2.75e^-7^; 0.5xK - 2.0xK: 2.55e^-6^, with p < 0.05/3, where 3 is the Bonferroni correction). D. Temperature-dependence of the channel from 5°C to 21°C (from black to red).

The amplitude of the change in input resistance depends on how much the temperature is altered (Fig. 3D), although the reversal potential is not altered. This suggests that larger changes in temperature activate this current to a greater extent. The behavior of this temperature-sensitive K^+^ current suggests it may be due to one or more of the 2-pore K^+^ channels such as TREK 1, TREK 2, TRAAK [13, 19, 24-27] that have been previously described.

This temperature-sensitive K^+^ current in stomach muscles is insensitive to the classical K^+^ channel antagonist, tetraethylammonium (TEA) [28, 29]. Bath application of 10^-2^M TEA did not change the reversal potential or the difference in current between the two temperatures seen at -65mV (Fig. 4A-C).

**Fig. 4.**
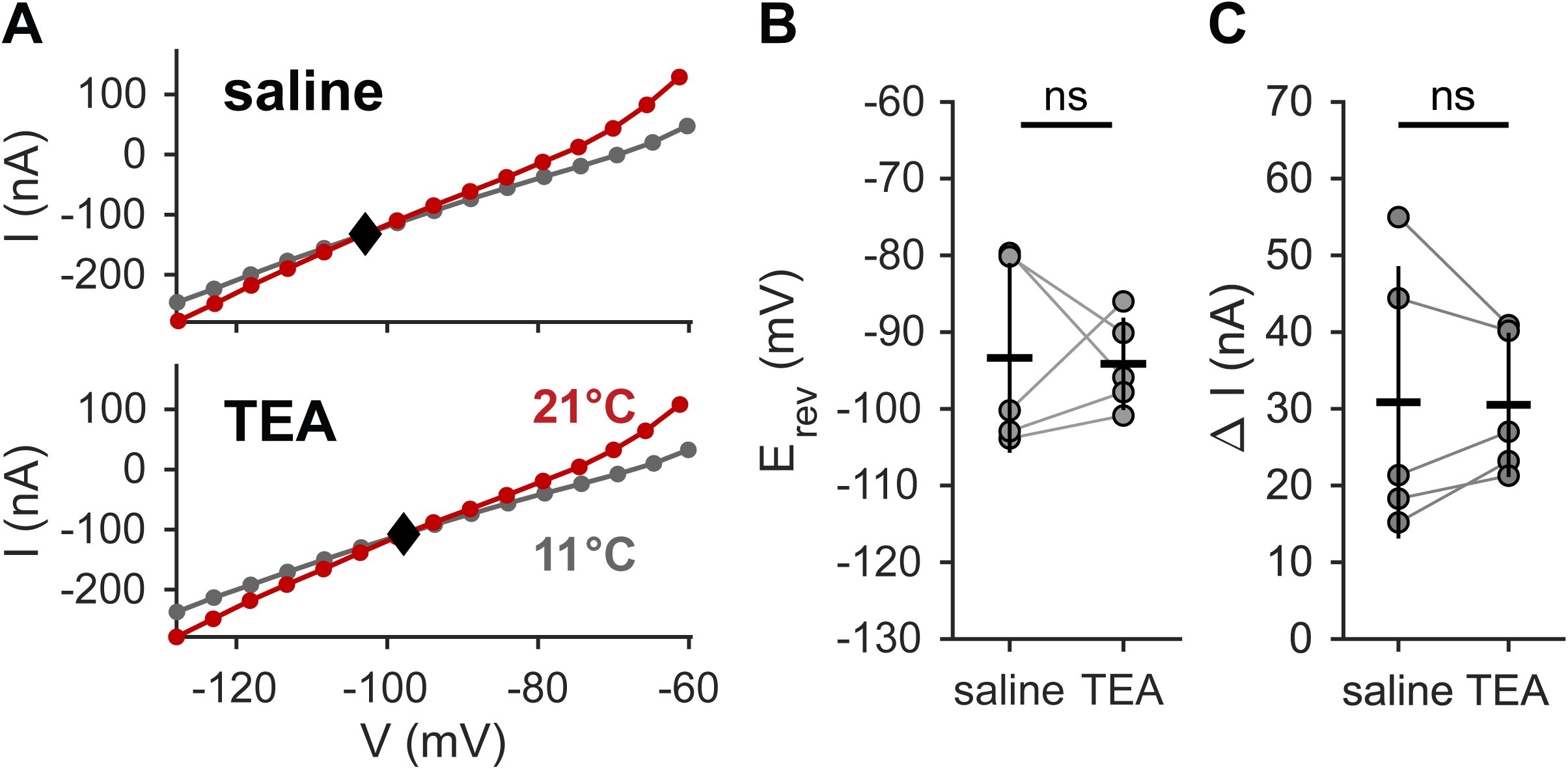
The temperature-dependent potassium channel is not sensitive to TEA in gm5b. A. IV curve at 11°C and 21°C in saline (top) and TEA (10^-2^ M, bottom), marked with the reversal potential (E_rev_). B. The reversal potential is not significantly different in saline and TEA. The horizontal and vertical lines represent the mean and standard deviation, respectively. Each dot represents a single muscle fiber. (Paired t-test, n = 5, p = 0.8973, p > 0.05). C. The difference in current (ΔI) obtained at 21°C and 11°C at -65 mV, compared between saline and TEA, shows no significant difference. The horizontal and vertical lines represent the mean and standard deviation. Each dot represents a single muscle fiber. (Paired t-test, n = 5, p = 0.9365).

Some 2-pore K+ channels (TASK1, 3, TREK 1,2, TWIK1, TRAAK) are characteristically pH sensitive [16, 17, 26, 30]. Consequently, we studied the effects of altered pH on EJPs (Fig. 5A) and EJCs (Fig. 5B). While both EJP and EJC amplitudes decrease with increasing temperature, their responses to pH were different. We observed a functional pH range in which activity remained stable (approximately pH 6.5–8.5). Deviations outside this range led to reduced amplitudes and, in some cases, abnormal activity such as spontaneous firing. A similar pH-dependent operating range has been described in stomatogastric ganglion neurons [31, 32].

**Fig. 5.**
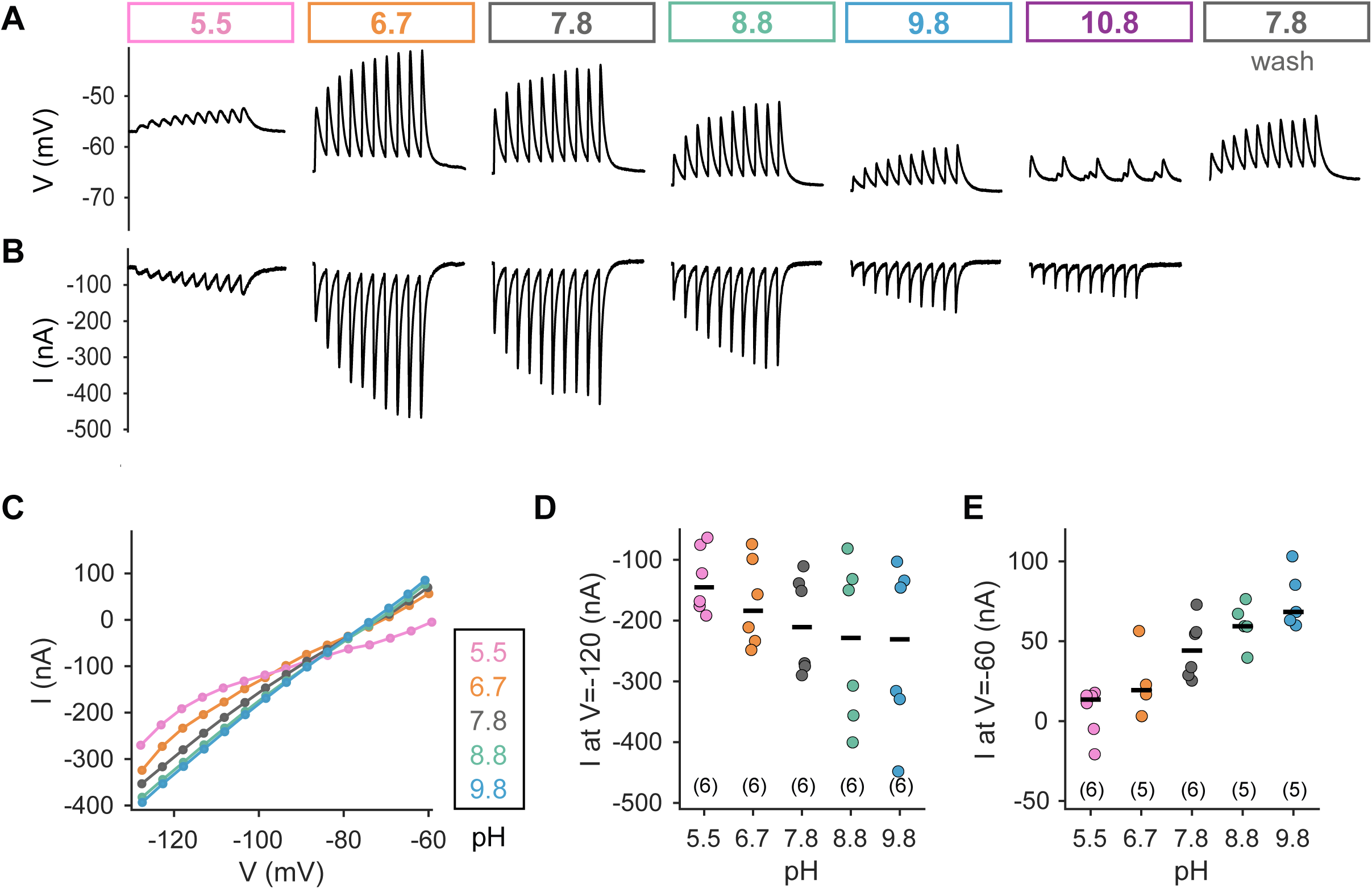
Effects of pH on EJPs in gm5b. A. Current-clamp recording of the muscle fiber while the lgn is stimulated at 5 Hz for 2 seconds in pH solutions ranging from 5.5 to 10.8. Both acidic and basic solutions affect the EJP amplitude, with optimal activity observed around the saline pH of 7.8. B. gm5b is clamped in a two-electrode voltage clamp at -80mV for the same stimulus. C. Voltage assessment of the pH sensitivity of the channel in gm5b. The IV curve of gm5b in different pH conditions shows that increasing pH activates the potassium channel. D-E. Current at -120 mV and -60 mV confirms the outward and inward activation by basic solutions. The horizontal and vertical lines represent the median and the standard deviation.

To directly assess pH sensitivity of this channel, we voltage-clamped muscle fibers in gm5b at different pH values and overlaid the resulting current traces (Fig. 5C). Increasing pH enhanced current activation, with an extrapolated reversal potential near –85 mV, once again suggesting a potassium conductance. Current measurements at –120 mV and –60 mV further confirmed the outward and inward currents elicited by basic solutions (Fig. 5D-E).

Two candidate 2 pore K^+^ channel genes were previously identified in *Cancer borealis* by sequence homology, referred to as *KCNK1* (Accession # KU681438) and *KCNK2* (KU681437). Positive expression of both *KCNK1* and *KCNK2* was detected in all three muscle types via qRT-PCR (Fig. 6). There were no significant quantitative differences detected among expression levels of either gene across muscle type.

**Fig. 6.**
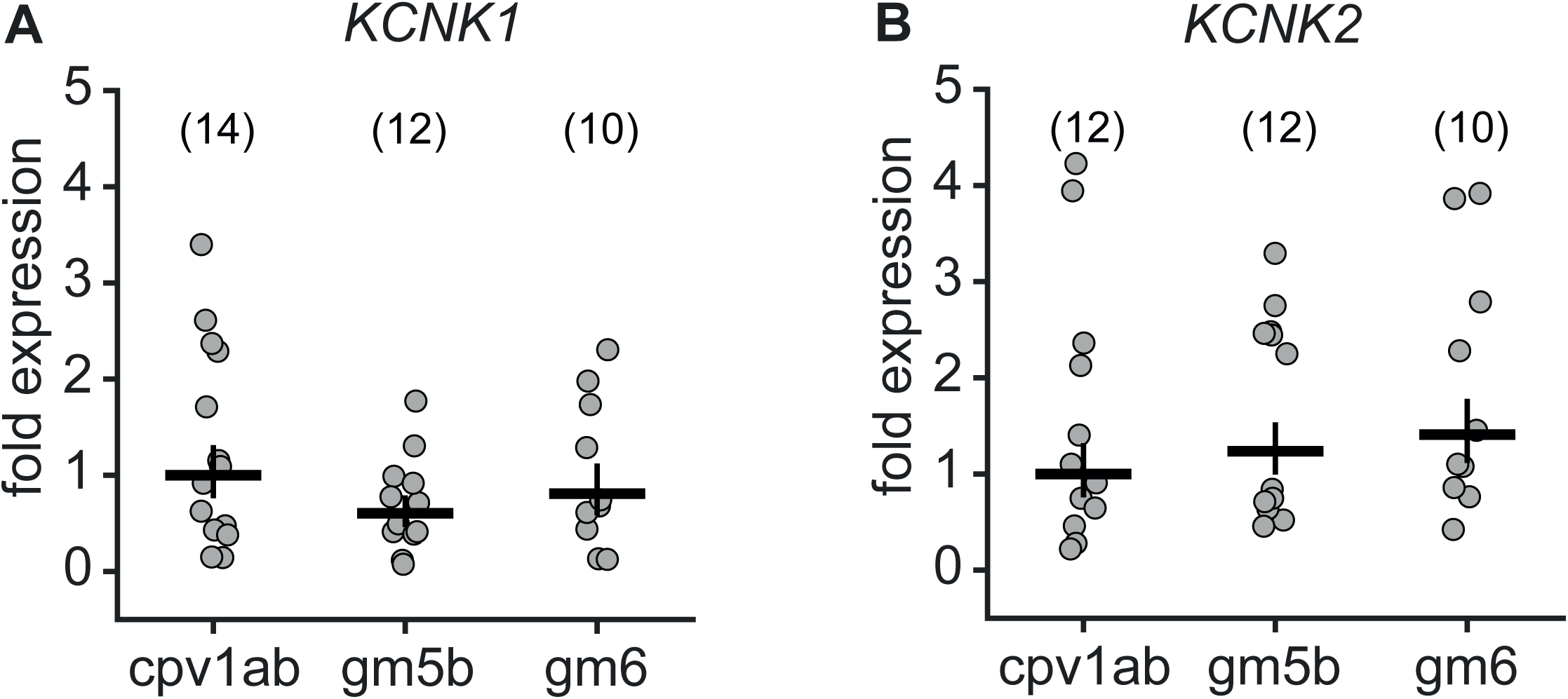
Steady-state mRNA levels of *KCNK1* and *KCNK2* in crab muscle. Relative mRNA levels are shown for the putative two-pore K^+^ channels *KCNK1* and *KCNK2* from cpv1ab, gm5b, and gm6 muscle from individual animals. Each data point represents one tissue sample from one animal. For both genes, each sample is expressed as a fold difference relative to the median Cq value of the cpv1ab group. The horizontal lines represent the mean of the fold expression, and the vertical lines span the lower and upper limits of the standard error. There were no significant differences across muscle types for either gene.

## Discussion

Many marine crustaceans, such as lobsters and crabs in the New England waters, encounter substantial temperature and pH fluctuations both daily and seasonally [1-4], and yet their motor systems remain robust within a substantial range. Most of the previous work on the effects of temperature and pH on the stomatogastric nervous system has been done on the isolated nervous system [31-39]. Nonetheless, the actual stomach movements produced by the animal depend both on the motor patterns and the properties of the neuromuscular synapses and the muscles themselves. Consequently, we now ask how temperature influences the neuromuscular junctions made by the stomatogastric ganglion neurons.

The effects of temperature on membrane potential and EJP amplitude of the crab stomach muscles (this paper) are consistent with earlier reports on many other crustacean muscles and neuromuscular junctions [9-12, 40, 41]. Given the substantial differences in ion channel expression and function across muscles and species, it is notable that temperature associated hyperpolarization of the muscle membrane potential appears to be a general property. Moreover, despite the differences in the synaptic dynamics and transmitters across different stomach muscles [6, 8], it is interesting that all stomach muscles display consistent hyperpolarizing responses to increases in temperature (Fig. 1). While these effects were historically attributed to a leak conductance or Na^+^/K^+^ pump activity [9-12], our results point to a more specific mechanism: the activation of potassium conductance mediated by two-pore domain K^+^ (K2P) channels.

Voltage-clamp evidence (Figs. 3 and 4) demonstrates that the reversal potential of the conductance affected by temperature is shifted by changes in K^+^ concentration, as would be expected for a K^+^ conductance. Transcriptomic data identified the expression of two putative K2P channel types in *Cancer borealis* [42], and Figure 6 confirms the presence of both of these transcripts in stomach muscles. When they were first identified by sequence homology, these channels were designated *Cb-KCNK1* and *Cb-KCNK2*, although unfortunately and confusingly, these designations do not correspond to the same numbered homologs in mammals: the *Cancer borealis KCNK1* coding sequence shares greatest homology with mammalian *KCNK9*/TASK-3 channels, while *Cancer borealis KCNK2* is most similar to mammalian *KCNK3*/TASK-1.

Within the mammalian two-pore potassium (K2P) channel family, TREK-1, TREK-2, and TRAAK (encoded by *KCNK2*, *KCNK10*, and *KCNK4*, respectively) are well-known polymodal sensors, activated by temperature, mechanical stretch, arachidonic acid, and volatile anesthetics. Our specific search of both the *Cancer borealis* genome [23] and neural transcriptome [42] to identify candidate homologs for *KCNK2, KCNK10* and *KCNK4* failed to detect such sequences in the crab. By contrast, TASK-1, TASK-3, and TWIK-1 (encoded by *KCNK3*, *KCNK9*, and *KCNK1*) are particularly sensitive to extracellular pH, and the identified crab *KCNK* channel sequences show clear homology to TASK-1 and TASK-3. Interestingly, both TREK-1 and TREK-2 display strong activation by temperature, while other K2P family members show more modest thermal sensitivity. An increase in pH activates TASK-1, TASK-3, TWIK-1, and TREK-1, whereas TREK-2 is instead activated by acidification [16, 30, 43, 44].

The crab genome appears to contain many fewer K2P channels than mammals, and we do not know whether both of the crab sequences identified encode channels that have both temperature and pH sensitivity, whether the two functions are separately conferred by the two genes, or some as of yet identified crab protein may be responsible for the thermal sensitivity we measured. The hyperpolarization with temperature and the decrease in EJP amplitude were monotonic across 11–25°C, whereas the response to pH was more complex. EJP amplitudes were maximal between pH 6.7 and 8.8, but declined outside this range, where abnormal activity such as spontaneous firing was also observed. Unlike the temperature response, the changes in resting membrane potential and input resistance with pH were not monotonic, suggesting that pH could act differently on the different ion channels. The existence of an optimal pH window is consistent with prior reports in pyloric and gastric neurons [31, 32, 35].

Taken together, these observations raise the possibility that in *Cancer borealis*, at least two K2P channel subtypes contribute to muscle responses. These channels may share similar temperature activation profiles but diverge in their pH sensitivity, providing a plausible explanation for the monotonic thermal response yet non-monotonic and process-dependent pH effects we observed.

We attempted to pharmacologically block the suspected K2P current using agents known to affect certain K2Ps in other systems [28]. In our experiments, unlike those in other systems, adding Ba^2+^ or quinine to the bath did not cleanly abolish the temperature-sensitive K^+^ current. Instead, these drugs induced side effects such as spontaneous muscle fiber firing and other conductance changes, which confounded the results. These side effects likely arise because Ba^2+^ or quinine may act at several neuronal and muscle targets. The natural genetic diversity expressed in the population of wild-caught animals used here may contribute as well.

Functionally, these channels provide a plausible mechanism for maintaining robust neuromuscular performance during environmental change. The pyloric and gastric rhythms accelerate 2–3-fold as temperature rises from 11°C to 21°C [33, 38, 45]. As the phase relationships and coupling across networks are preserved [38], we see an increase in the spike frequency within the burst – acting as a direct input to the neuromuscular junction. Without compensation, a depolarized muscle receiving inputs at twice the spike frequency would contract more strongly or continuously, risking abnormal movement pattern. By hyperpolarizing fibers as temperature increases, K2Ps reduce excitability and EJP amplitude, partially offsetting the enhanced synaptic drive. Thus, muscle fibers themselves contribute to temperature compensation, complementing circuit-level robustness described previously in the STG [33, 34, 46].

In conclusion, we provide evidence that temperature- and pH-sensitive K2P channels contribute to the modulation of muscle excitability under varying environmental conditions. These findings revise our understanding of how crustacean muscles adapt physiologically to temperature and pH fluctuations. We propose that these channels play a critical role in maintaining the appropriate amplitude and timing of muscle contractions as motor pattern frequency changes with temperature. More broadly, this work opens new avenues for exploring temperature compensation at the neuromuscular junction and how multiple environmental variables may affect the motor function in ectothermic animals.

## Star Methods

### Detailed methods are provided in the paper and include the following

- Key resource table
- Resource availability

- Lead contact
- Material availability
- Data and code availability
- Experimental model and subject details

- Experimental preparation
- Electrophysiology
- Reverse Transcription and Real-Time Quantitative Polymerase Chain Reactions
- Quantification and statistical analysis

## Acknowledgments

This work was supported by R35 NS 097343 (EM), Belgian American Education Foundation – BAEF, Wallonia-Belgium International – WBI Postdoctoral Fellowship (KJ).

We thank Sonal Kedia and Kyra Schapiro for their support and valuable feedback on this project, Margaret Lee and Maria Ivanova for training assistance, Jackie Seddon for help with the installation of the muscle recording rig, and Gwen Harris for management of the animals and lab facility.

## STAR METHODS

### KEY RESOURCE TABLE

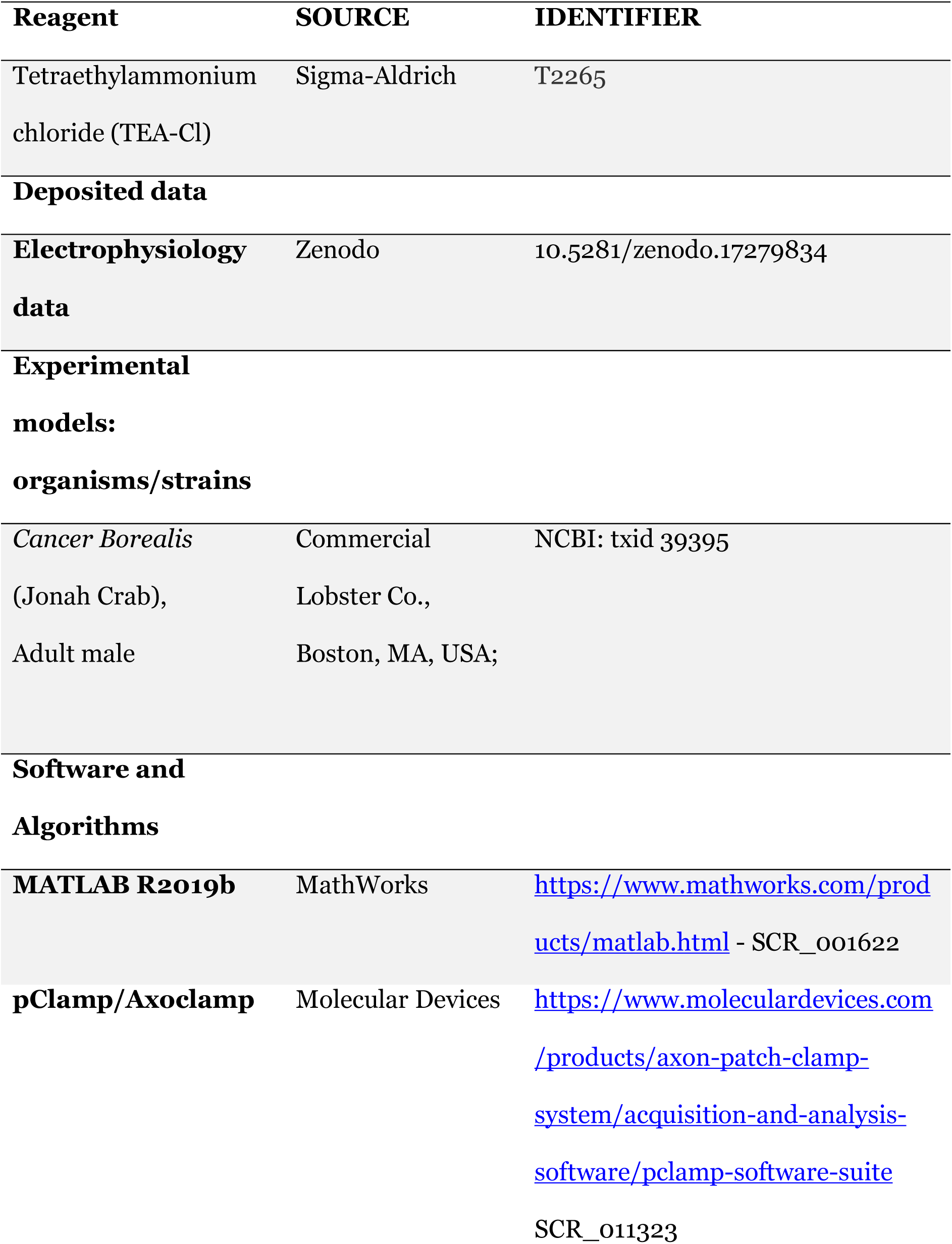
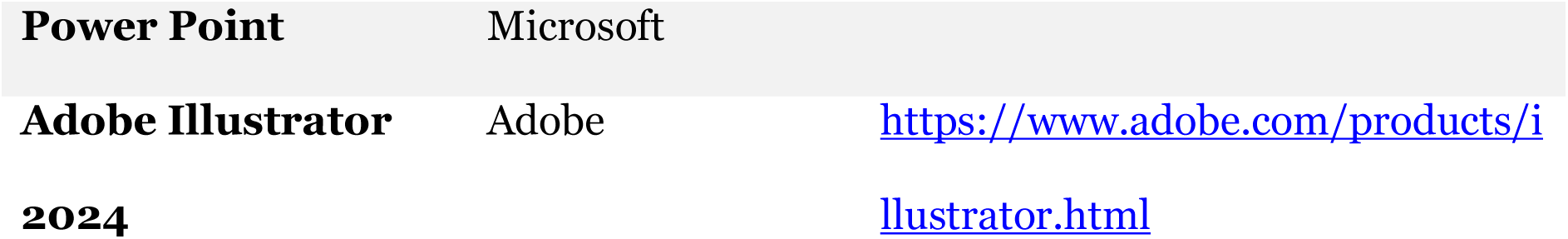

### RESOURCE AVAILABILITY

#### Lead contact

Further information and requests for resources and reagents should be directed to and will be fulfilled by the Lead Contact, Kathleen Jacquerie.

#### Material availability

This study did not generate any new unique reagents.

#### Data and code availability

All electrophysiology data and metadata used for this study can be found at 10.5281/zenodo.17279834. Custom MATLAB codes extracting the resting membrane voltage, the IV curves steady-state data, or the plotting functions are available on demand.

### EXPERIMENTAL MODEL AND SUBJECT DETAILS

Adult male Jonah Crabs (*Cancer borealis*) were purchased from Commercial Lobster Company (Boston, MA) and held in filtered, circulating, aerated, artificial seawater (Instant Ocean Sea Salt Mix) at 11°C. Crabs were acquired between May 2024 and June 2025. Animals were typically held in the laboratory for one week before use.

### METHOD DETAILS

#### Experimental preparation

Stomach musculature and the stomatogastric nervous system (STN) were dissected following the description and nomenclature of [5]. Nerve dissection followed the procedures described by [48]. Specific neuromuscular preparations were isolated from the foregut. Before starting experiments, the fat layer overlaying the muscles was carefully removed to improve access for recording from muscle fibers while keeping nerve terminals attached [49, 50].

Reduced preparations of parts of the stomach and nerves were pinned into a silicone elastomer-lined (Sylgard 184: Dow Corning) dish. The preparation was maintained in *Cancer borealis* saline solution (440 mM NaCl, 11 mM KCl, 26 mM MgCl2, 13 mM CaCl2, 11 mM Trizma base, 5.4mM maleic acid, pH 7.8 at 11°C) throughout the dissection and experiment. Saline was continuously perfused at 8-11 mL/min via a peristaltic pump (model Ismatec Ecoline). Temperature was adjusted from 5°C to 21°C using a Peltier system (Warner Instruments SC-20) with a temperature controller (CL-100, Warner Instruments) and a thermocouple probe in the bath near the recording site. Temperatures within ±0.3°C of the target value (e.g., 10.7–11.3°C for 11°C) were considered as 11°C for analysis.

High K^+^ saline (2xK^+^, 22mM KCl) and low K^+^ saline (0.5xK^+^, 5.5mM) were prepared by adding more KCl salt to the normal saline. TEA-containing saline was prepared by adding 10^-2^M of tetraethylammonium chloride (Sigma Aldrich). Additional quantities of Trizma base or maleic acid were added to achieve solutions with pH 5.5, 6.5, 7.5, 8.5, 9.5 at room temperature and at 11°C pH 5.5, 6.7, 7.8, 8.8, 9.8, and 10.8 according to (Haley et al, 2018). For a more basic solution, NaOH was used to reach 10.5 at room temperature. Solutions were calibrated using the device Mettler Tolledo pH meter. Values within ±0.2 pH units of the target pH (e.g., 6.4–6.6 for pH 6.5) were considered as pH 6.5 for analysis.

#### Electrophysiology

Electrophysiology experiments were performed as previously described [49-51]. Muscles were pinned to Sylgard dishes. Nerves were stimulated using suction electrodes attached to the cut nerve end, connected to an AM Systems isolated pulse stimulator (Model 2100). Intracellular recordings of excitatory junctional potentials (EJPs) in muscle fibers were obtained using sharp borosilicate glass microelectrodes (Sutter Instrument; ID 0.86 mm, OD 1.5 mm), pulled with a micropipette puller (Sutter Instrument, P-97) and backfilled with 400mM K_2_S0_4_, 20mM KCl, 33mM NaCl. Electrode resistances were between 5 and 20MΩ.

Electrodes were mounted on Leica Leitz mechanical micromanipulators with HS-2A-x1LU Axoclamp 2B and HS-9A-x1U Axoclamp 900A headstages (Molecular Devices). Signals were amplified using Axoclamp 2B or 900A amplifiers (Molecular Devices), digitized at 10 kHz using a Digidata 1440, and recorded in Clampex 10.7. The electrodes can have an offset between 0 and 3mV for a 10°C increase from 11°C to 21°C. Pulse trains (2 s duration) were delivered at 2 or 5 Hz. Individual pulse durations ranged from 0.2 to 0.9 ms. Pulse amplitudes were calibrated to reliably trigger single EJPs at 11°C while keeping a 10–20% safety margin. Muscle fibers depolarized beyond −50 mV were excluded from recordings [52].

Excitatory junctional currents (EJCs) were recorded in two-electrode voltage clamp mode at −80 mV. Two electrodes were inserted in the same fiber along the longitudinal axis. The distance between the two electrodes was less than the diameter of the fiber. Current-voltage (I-V) curves were obtained by stepping the voltage from −130 to −60 mV (starting from −80 mV) for 2 s. Saline levels were minimized to reduce capacitive coupling. The headstage for recording currents was a Molecular Devices HS-2A-x10MU Axoclamp 2B, connected via an adapter to the Axoclamp 900A. Because the muscle fibers are large and electrically coupled, producing substantial currents, voltage steps were accepted only if they reached the intended values and the recorded currents remained stable.

#### Reverse Transcription and Real-Time Quantitative Polymerase Chain Reactions (qRT-PCR)

After electrophysiology experiments, whole muscles from the left and right sides of each animal were collected and stored at −80 °C. The cpv1a and cpv1b muscles were pooled and analyzed together (hereafter, “cpv1ab”). Tissues were homogenized in 750μl TRIzol (Invitrogen) for RNA extraction. Total RNA was isolated according to the manufacturer’s protocol (Invitrogen). 100 ng of total RNA from each sample was reverse transcribed using qScript reverse transcriptase (QuantaBio) primed with a mixture of oligo-dT and random hexamers. The final volume of the reverse transcription reaction was 20 μl and contained a final concentration of 2.5 ng/μl random hexamers, 2.5 μM oligo-dT, 40 U of RNase inhibitor, and 200 U of reverse transcriptase. Following heat inactivation of the enzyme, samples were diluted in ultrapure water to a final volume of 50 μl before this template was used in qRT-PCR analyses.

cDNA was used in qRT-PCR reactions for *KCNK1* and *KCNK2* genes with primer sets previously validated in this system [53]. qRT-PCR reactions consisted of primer pairs at a final concentration of 2.5 μM, cDNA template (equivalent to 3.33 ng of total cellular RNA), and SsoAdvanced SYBR mastermix (BioRad) according to the manufacturer’s instructions. Reactions were carried out on a CFXConnect (BioRad) machine with a three-step cycle of 95°C-15s, 58°C-20s, 72°C-20s, followed by a melt curve ramp from 65°C to 95°C. Data were acquired during the 72°C step and every 0.5°C of the melt curve. All reactions were run in triplicates of 10 μl, and the average Cq (quantification cycle) was used for analysis. Reactions were normalized to a fixed amount of total cellular RNA [54, 55] and expression level for each individual sample was then calculated as a fold-expression level relative to the median Cq of the cpv1ab muscle group as follows: *Fold Expression_x_* = 2^−(*Cq_x_− Cq_medianCtrl_*)^.

### QUANTIFICATION AND STATISTICAL ANALYSIS

We developed custom MATLAB scripts to visualize and analyze recordings obtained using Clampex software. These scripts were used to extract the resting membrane potential, the IV curve, the input resistance (R_in_), and the reversal potential (E_rev_).

Resting membrane potential values were obtained by averaging over 0.5 seconds prior to the onset of the pulse train. Currents associated with each voltage step were averaged over the final 0.17 seconds of the 2-second voltage steps to ensure steady-state measurements. Input resistance was calculated between −92 mV and −73 mV to remain within the linear portion of the IV curve using MATLAB’s fit function with a first-degree polynomial. Reversal potential was determined by calculating the intersection of the IV curves at 11°C and 21°C. Raw traces were plotted and smoothed using 8–30 points (MATLAB smooth function), while voltage-clamp current traces were smoothed using ∼10 points.

For statistical analyses, data are reported as either the median or the mean ± standard deviation (SD), as indicated in the figure legends. Paired t-tests were used to compare the same preparation across different conditions. Bonferroni corrections were applied to adjust for multiple comparisons using the Wilcoxon rank-sum test. Significant differences are indicated as follows: *p < 0.05, **p < 0.01, ***p < 0.001, ****p < 0.0001. P-values lower than 1e⁻⁶ are reported as 0.00000.

