## Supplementary Information for "Temperature and pH-dependent Potassium Currents of Muscles of the Stomatogastric Nervous System of the Crab, *Cancer borealis*"

**Table S1 – Statistical features used in the different figures.** n represents the number of muscle fibers, Temp=Temperature, Med=Median, SD=Standard Deviation, CV=coefficient of variation, N.A.=Not available, cpv1ab represent the extraction of cpv1a and cpv1b. For Fig. 6, STD and CV are replaced by the intervals given by the lower and upper limits of the fold expression.

| Fig | Measure | Muscle | Solution | Temp | n | Mean | Med | SD | CV |
|---|---|---|---|---|---|---|---|---|---|
| 2A | $V_{rest}$ (mV) | gm5b | saline | 11°C | 68 | -69.92 | -69.53 | 4.600 | 0.06579 |
| 2A | $V_{rest}$ (mV) | gm5b | saline | 21°C | 68 | -77.94 | -77.67 | 4.556 | 0.05845 |
| 2A | $V_{rest}$ (mV) | gm6 | saline | 11°C | 48 | -69.71 | -69.40 | 4.470 | 0.06413 |
| 2A | $V_{rest}$ (mV) | gm6 | saline | 21°C | 48 | -79.10 | -78.93 | 4.268 | 0.05396 |
| 2A | $V_{rest}$ (mV) | gm8a | saline | 11°C | 12 | -71.48 | -72.18 | 3.758 | 0.05258 |
| 2A | $V_{rest}$ (mV) | gm8a | saline | 21°C | 12 | -79.67 | -79.83 | 3.804 | 0.04774 |
| 2A | $V_{rest}$ (mV) | cpv1a | saline | 11°C | 24 | -63.91 | -63.71 | 4.469 | 0.06992 |
| 2A | $V_{rest}$ (mV) | cpv1a | saline | 21°C | 24 | -73.44 | -73.65 | 5.138 | 0.06996 |
| 2A | $V_{rest}$ (mV) | cpv4 | saline | 11°C | 4 | -67.49 | -66.64 | 4.131 | 0.06121 |
| 2A | $V_{rest}$ (mV) | cpv4 | saline | 21°C | 4 | -76.62 | -74.78 | 6.945 | 0.09064 |
| 2A | $V_{rest}$ (mV) | cpv6 | saline | 11°C | 5 | -68.86 | -67.30 | 9.858 | 0.1432 |
| 2A | $V_{rest}$ (mV) | cpv6 | saline | 21°C | 5 | -77.62 | -80.80 | 8.592 | 0.1107 |
| 2B | $R_{in}$ (mΩ) | gm5b | saline | 11°C | 21 | 339.0 | 266.9 | 218.2 | 0.6437 |
| 2B | $R_{in}$ (mΩ) | gm5b | saline | 21°C | 21 | 222.0 | 169.6 | 147.2 | 0.6633 |
| 2B | $R_{in}$ (mΩ) | gm6 | saline | 11°C | 8 | 153.7 | 144.7 | 30.00 | 0.1952 |
| 2B | $R_{in}$ (mΩ) | gm6 | saline | 21°C | 8 | 94.92 | 83.25 | 25.01 | 0.2634 |
| 2B | $R_{in}$ (mΩ) | cpv1a | saline | 11°C | 6 | 367.7 | 369.7 | 127.5 | 0.3468 |
| 2B | $R_{in}$ (mΩ) | cpv1a | saline | 21°C | 6 | 244.2 | 272.8 | 106.3 | 0.4352 |
| 3C | $E_{rev}$ (mV) | gm5b | 0.5xK | - | 17 | -107.1 | -106.9 | 8.989 | 0.0839 |
| 3C | $E_{rev}$ (mV) | gm5b | 1.0xK | - | 37 | -96.58 | -96.53 | 12.71 | 0.1316 |
| 3C | $E_{rev}$ (mV) | gm5b | 2.0xK | - | 14 | -72.12 | -71.45 | 6.593 | 0.09142 |
| 4B | $E_{rev}$ (mV) | gm5b | saline | - | 5 | -93.36 | -100.2 | 12.35 | 0.1323 |
| 4B | $E_{rev}$ (mV) | gm5b | TEA $10^{-2}$M | - | 5 | -94.14 | -95.89 | 6.012 | 0.06386 |
| 4C | ΔI (nA) | gm5b | saline | - | 5 | 30.85 | 21.36 | 17.74 | 0.5752 |
| 4C | ΔI (nA) | gm5b | TEA $10^{-2}$M | - | 5 | 30.51 | 27.01 | 9.938 | 0.3076 |
| 5D | I (nA) | gm5b | pH 5.5 | 11°C | 6 | -132.8 | -145.2 | 54.38 | 0.4095 |
| 5D | I (nA) | gm5b | pH 6.7 | 11°C | 6 | -170.4 | -183.9 | 72.55 | 0.4259 |
| 5D | I (nA) | gm5b | pH 7.8 | 11°C | 6 | -205.9 | -210.7 | 80.74 | 0.3921 |
| 5D | I (nA) | gm5b | pH 8.8 | 11°C | 6 | -237.7 | -228.5 | 133.2 | 0.5602 |
| 5D | I (nA) | gm5b | pH 9.8 | 11°C | 6 | -246.0 | -230.8 | 138.3 | 0.5623 |
| 5E | I (nA) | gm5b | pH 5.5 | 11°C | 6 | 5.846 | 13.51 | 15.45 | 2.642 |
| 5E | I (nA) | gm5b | pH 6.7 | 11°C | 5 | 23.70 | 19.36 | 19.75 | 0.8335 |
| 5E | I (nA) | gm5b | pH 7.8 | 11°C | 6 | 45.21 | 44.19 | 18.76 | 0.4147 |
| 5E | I (nA) | gm5b | pH 8.8 | 11°C | 5 | 60.39 | 59.42 | 13.53 | 0.224 |
| 5E | I (nA) | gm5b | pH 9.8 | 11°C | 5 | 76.01 | 68.33 | 18.07 | 0.2377 |
| 6A | Fold diff (-) | cpv1ab | saline | - | 14 | 1.000 | 1.000 | [0.7611; 1.314] | |
| 6A | Fold diff (-) | gm5b | saline | - | 12 | 0.604 | 0.5946 | [0.4614; 0.7909] | |
| 6A | Fold diff (-) | gm6 | saline | - | 10 | 0.808 | 0.7120 | [0.5824; 1.122] | |
| 6B | Fold diff (-) | cpv1ab | saline | - | 12 | 1.000 | 1.000 | [0.758; 1.320] | |
| 6B | Fold diff (-) | gm5b | saline | - | 12 | 1.237 | 1.3755 | [0.995; 1.537] | |
| 6B | Fold diff (-) | gm6 | saline | - | 10 | 1.410 | 1.2658 | [1.115; 1.782] | |
